# Cdc42 promotes epithelial morphogenesis by coupling Par-complex and Crumbs recruitment via Par6-aPKC

**DOI:** 10.1101/513028

**Authors:** Francisca Nunes de Almeida, Rhian F. Walther, Mary Pressé, Evi Vlassaks, Franck Pichaud

**Author notes:** Co-first authors (Francisca Nunes de Almeida, Rhian F. Walther).

## Abstract

Epithelial morphogenesis depends on Cdc42, which acts in part through the Par-complex (Baz/Par3-Par6-aPKC), and Crumbs (Crb/CRB3). However, how these proteins come together and interact during morphogenesis is not well understood. Here we show that in *Drosophila* cells, Cdc42 is active at the apical membrane and cell-cell contacts, and couples apical recruitment of Par6-aPKC and Crb to promote apical identify. This coupling depends on Cdc42 and aPKC binding to Par6, and is mediated by Par6 binding to Crb. Conversely, Crb promotes the apical retention of Par6-aPKC, and thus Cdc42-Par6-aPKC and Crb promote each other’s apical accumulation to specify the apical membrane and *Zonula Adherens (ZA)*. Altogether, our work shows that epithelial morphogenesis relies on a two tiered-mechanism whereby Cdc42 coordinates Par-complex assembly and apical recruitment of Crb through Par6-aPKC.

## INTRODUCTION

Epithelial cell morphogenesis requires the specification and maturation of distinct plasma membrane domains along the apical (top) to basal (bottom) axis of the cell. These domains support specific functions and consist of the apical membrane, which contains microvilli flanked by sub-apical membranes, the lateral membrane, containing junctional domains that mediate cell-cell adhesion and function as paracellular barriers, and the basal membrane. Morphogenesis of these membrane domains depends on a set of proteins that are conserved through evolution. These include the apical *Par*titioning defective proteins Bazooka (Baz; Par3 in mammals) and Par6, the serine/threonine kinase aPKC (PKCζ/ɩ in mammals), the small GTPase Cdc42 (Goldstein and Macara, 2007; Hoege and Hyman, 2013; Joberty et al., 2000; Johansson et al., 2000; Kemphues, 2000; Lin et al., 2000), the transmembrane protein Crb (CRB3 in mammals) and its binding partner Stardust (Sdt; PALS1 in mammals) (Flores-Benitez and Knust, 2016; Tepass, 2012). However, how exactly these proteins come together to promote epithelial morphogenesis is not well understood.

*Drosophila* is a powerful model system to dissect the mechanism of epithelial polarity and morphogenesis. In *Drosophila* cells, the *Zonula Adherens (ZA)*, which contains Ecadherin (Ecad), marks the boundary between the apical and lateral membrane, and mediates cell-cell adhesion. Amongst the popular model epithelial cells to study the mechanisms of epithelial morphogenesis *in vivo* is the developing pupal photoreceptor. As it matures, this cell presents well-resolved microvilli, sub-apical and *ZA* domains. Morphogenesis of these three membrane domains requires the function of Cdc42, Baz, Par6, aPKC and Crb (Hong et al., 2003; Izaddoost et al., 2002; Muschalik and Knust, 2011; Nam and Choi, 2003; Pellikka et al., 2002; Walther and Pichaud, 2010). In addition, apical membrane and *ZA* morphogenesis in these cells requires the function of the Cdc42 effector kinases P21-activated kinase Pak4 and Ghenghis Khan/MRCK *(gek)*, which also regulate junction maturation and apical membrane morphogenesis in vertebrate cells (Jin et al., 2015; Schneeberger and Raabe, 2003; Wallace et al., 2010; Walther et al., 2016; Zihni et al., 2017).

Amongst the epithelial protein network, Baz/Par3 plays a key role during pupal photoreceptor morphogenesis by promoting the recruitment Par6, aPKC and Crb to the apical membrane (Hong et al., 2003; Walther et al., 2016; Walther and Pichaud, 2010). However, Baz localization does not depend on Par6, aPKC and Crb, which indicates that in these cells, Baz acts upstream of the regulatory network that promotes polarized morphogenesis. Similarly, in the early fly embryo, early apical localization of Baz does not depend on aPKC or Par6 (Harris and Peifer, 2005). The situation is somewhat different in the follicular epithelium, where Baz is dispensable for Par6, aPKC and Crb apical localization (Shahab et al., 2015), thus pointing to cell-type specific differences in the manner in which these proteins interact with each other during morphogenesis. A predominant role for Par3 during epithelial morphogenesis is conserved through evolution. In MDCK spheroids, Par3 is one of the earliest proteins that marks the apical domain during morphogenesis and is required for lumen formation (Bryant et al., 2010).

Next to Baz, Cdc42 is also required for the membrane localization of Par6, aPKC and Crb in the pupal photoreceptor (Walther and Pichaud, 2010). This requirement is likely to stem, at least in part, from the ability of Cdc42 to bind to Par6, which has been shown to be essential for Par6 recruitment at the apical membrane in the fly embryo (Hutterer et al., 2004). In addition, evidence is that Cdc42 binding to Par6 is required for Crb apical accumulation, as for example in the pupal photoreceptor (Walther and Pichaud). Par6 can bind to Crb/CRB3 (Lemmers et al., 2004), and the Par6 binding partner aPKC has been proposed to act downstream of Cdc42 to promote Crb accumulation at the apical membrane (Harris and Tepass, 2008). One possible mechanisms explaining how Par6-aPKC might promotes Crb recruitment, is that Crb is stabilized upon aPKC phosphorylation (Sotillos et al., 2004). However, preventing Crb phosphorylation *in vivo* does not affect epithelial polarity or morphogenesis (Cao et al., 2017), indicating that Par6-aPKC can regulate Crb stability through other routes.

Understanding the regulations that take place between Cdc42, Baz, Par6-aPKC and Crb is important because they underpin the morphogenesis of the apical membrane and ZA. For instance, binding of Par6 to Crb has been shown to outcompete Baz binding to Par6-aPKC (Morais-de-Sa et al., 2010), resulting in the exclusion of Baz from the Par-(Baz-Par6-aPKC) complex (Krahn et al., 2010; Morais-de-Sa et al., 2010; Walther and Pichaud, 2010). This exclusion mechanism acts in concert with phosphorylation of Baz by aPKC at a conserved Serine residue S980 (S827 in vertebrate Par3D) (Krahn et al., 2010; Morais-de-Sa et al., 2010; Nagai-Tamai et al., 2002; Walther and Pichaud, 2010) to limit Baz localization to the border between the apical and lateral membrane, where it is thought to promote *ZA* morphogenesis. In addition, Baz phosphorylation by aPKC promotes the release of Sdt from Baz, presumably to allow for Sdt to interact with Crb during apical membrane morphogenesis (Krahn et al., 2010). Therefore, Par6 binding to Crb is thought to act as a main regulation that drives the differentiation of the apical membrane and ZA. Interestingly, Cdc42 is thought to promote this binding (Kempkens et al., 2006; Lemmers et al., 2004; Whitney et al., 2016), but the functional relevance for this regulation is not fully understood.

Altogether, when considering the mechanisms of epithelial morphogenesis, Par6 occupies an interesting position within the epithelial polarity network because it can bind to Cdc42, Baz, aPKC and Crb (Supplementary Figure 1). This makes this PDZ-protein an ideal candidate to act downstream of Cdc42 to coordinate the function of the Par-complex and Crb during epithelial morphogenesis. To test this possibility, we made use of the pupal photoreceptor to perform an *in vivo* structure-function study of Par6 with a focus on disrupting its binding to Cdc42, aPKC and Crb. We complemented this work using the follicular epithelium, where Par6 function has not been extensively studied so far. Altogether our work shows that in a cell undergoing morphogenesis, Cdc42 is active at the apical membrane and ZA, and coordinates Par-complex assembly and Crb recruitment. Our results indicate that while Baz is required to load Par6-aPKC onto the apical membrane, Cdc42 binding to Par6 regulates the capture of Par6-aPKC by Crb. This regulation leads to the apical accumulation of Crb and relative depletion of Par6-aPKC from the ZA. In addition, we show that Crb is dispensable for the recruitment of an apical Par6-aPKC pool that separates from Baz, and our results further suggest that this pool of Par6-aPKC is bound to Cdc42.

## RESULTS

### Cdc42-GTP localizes at the apical membrane and *ZA*

In the early (37-40% APF) pupal photoreceptor (Figure 1A), the apical membrane consists of poorly differentiated ruffles and is not yet subdivided into apical microvilli and sub-apical membrane (stalk membrane) (Ready, 2002). Similarly, in the follicular epithelium, the apical membrane consists of microvilli with a very short sub-apical membrane apical and *ZA* (Tanentzapf et al., 2000) (Figure 1A). In both cell types, Crb and Par6-aPKC are enriched at the apical membrane while Baz and Arm are enriched at the *ZA* (Figure 1B-H) (Abdelilah-Seyfried et al., 2003; Franz and Riechmann, 2010; Hong et al., 2003; Izaddoost et al., 2002; Morais-de-Sa et al., 2010; Muschalik and Knust, 2011; Nam and Choi, 2003; Pellikka et al., 2002; Walther and Pichaud, 2010). Amongst these proteins, Cdc42 is required for the recruitment of aPKC and Crb at the apical membrane of the pupal photoreceptor (Figure 1I-J) and (Walther and Pichaud, 2010). One possibility is that Cdc42 is active and accumulates at the apical membrane and/or *ZA* to promote Par6-aPKC and Crb recruitment. To test this possibility, we monitored Cdc42 using a functional Cdc42::mCherry fusion protein (Abreu-Blanco et al., 2014). mCherry::Cdc42 expressed in *cdc42*^3^ and *cdc42*^4^ mutants rescues embryonic lethality and supports viability to eclosion, albeit at less than the expected Mendelian ratio. We found that mCherry::Cdc42 accumulates at the apical membrane of the developing pupal photoreceptor and follicular epithelium, and is also present at the *ZA* (Figure1K-L). To complement this analysis, we generated a GFP probe (WASp-CRIB::GFP) that binds to active, GTP loaded Cdc42 (Figure 1M). In pupal photoreceptors, WASp-CRIB::GFP showed a cytosolic staining and an enrichment at the apical membrane as well as the *ZA* (Figure 1N). A mutant version of the probe that cannot bind GTP-Cdc42 (WASp-CRIB::GFP^MUT^) (Figure 1M), showed a cytosolic staining similar to that detected with WASp-CRIB::GFP, but lacked the membrane staining (Figure 1O), demonstrating that the apical and *ZA* staining obtained with WASp-CRIB::GFP is specific to Cdc42-GTP. Therefore, we conclude that Cdc42 is present and active at the apical membrane and *ZA* of the developing pupal photoreceptor.

**Figure 1:**
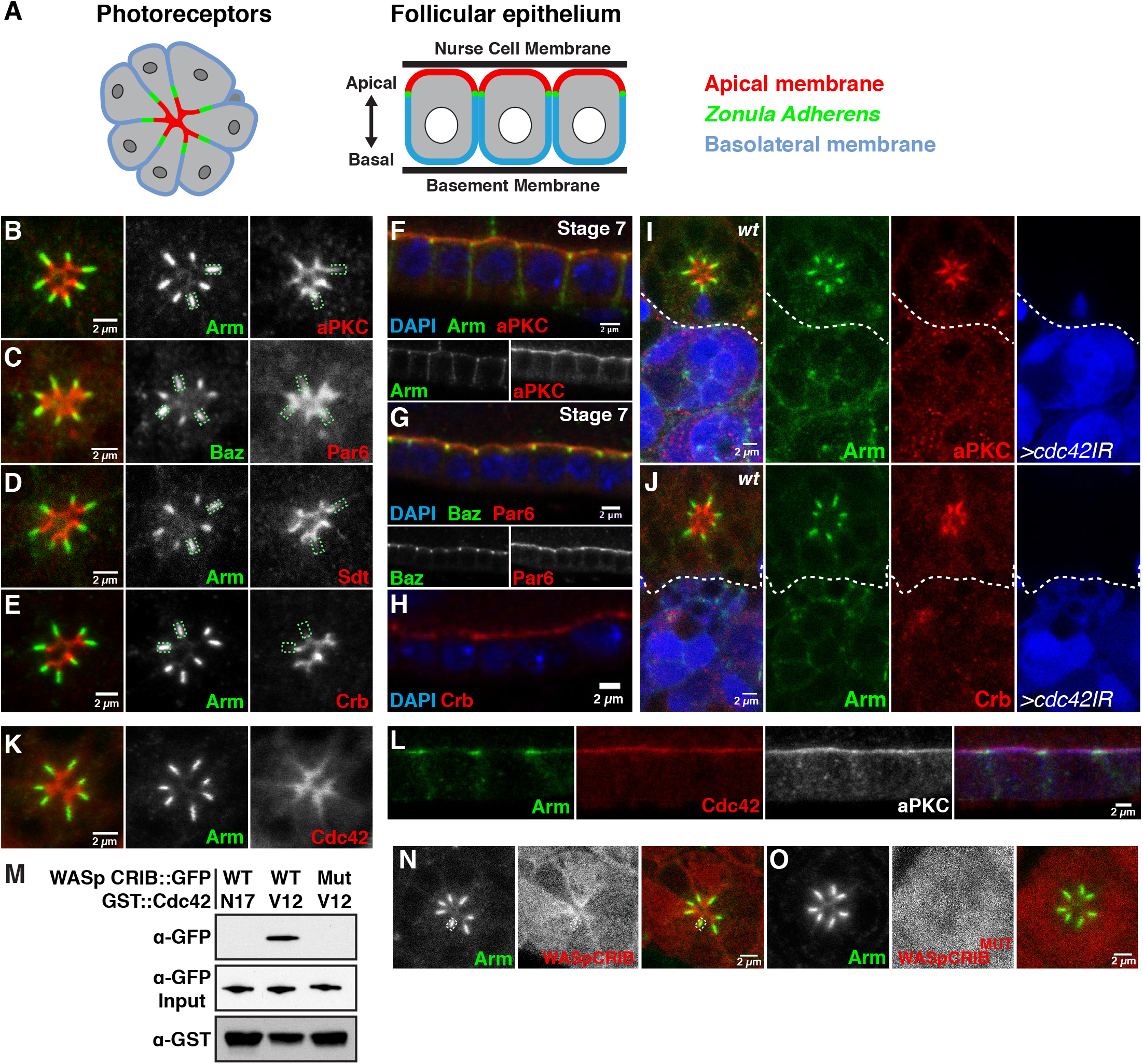
Cdc42-GTP localizes at the apical membrane and *ZA*. (**A**) Schematic representation of the developing fly photoreceptors at early-pupal stage, and stage 7 follicular epithelial cells. Photoreceptors are arranged in a circular cluster called an ommatidium. The photoreceptor apical membranes (red) face the central lumen of the cluster (*ZA*, green; basolateral, blue). (**B-E**) Early-pupal photoreceptors stained for (**B**) Arm (green) and aPKC (red), (**C**) Baz (green) and Par6 (red), (**D**) Arm (green) and Sdt (red), and (**E**) Arm (green) and Crb (red). Green rectangles delineate the *ZA* to show the relative overlap between Arm/Baz and aPKC/Par6/Sdt/Crb. (**F-H**) Wild-type cuboidal follicle cells from stage 7 follicles stained for (**F**) DAPI (blue), Arm (green) and aPKC (red), (**G**) DAPI (blue), Baz (green) and Par6 (red), and (**H**) DAPI (blue) and Crb (red). (**I**) Cdc42IR marked by the presence of GFP (blue) and stained for Arm (green), aPKC (red) and (**J**) Arm (green), Crb (red). (**K**) mCherry::Cdc42 (red) and Arm (green) localization in the pupal photoreceptor. (**L**) mCherry::Cdc42 (red), Arm (green) and aPKC (grey; blue in merged panel) localization in the follicular epithelium, stage 7. (**M**) Representative pull down experiment combining GST::Cdc42V12, GST::Cdc42N17 and the Cdc42-GTP probe WASp-CRIB::GFP or its mutated version (WASp-CRIB::GFP^MUT^) expressed in S2R+ cells. (**N**) Localization of Cdc42-GTP monitored using WASp-CRIB::GFP (red), Arm (green). A white rectangle delineates a ZA to show that WASp-CRIB::GFP can be detected in this membrane domain. (**O**) Control WASp-CRIB::GFP^MUT^ (red), Arm (green). All scale bars = 2 microns.

### Par6 is required for apical recruitment of aPKC and Crb

*par6* is required for cell viability in the retina, (Walther and Pichaud, 2010), however by raising the animals at 18 degrees we could recover photoreceptors mutant for the null allele *par6^Δ226^*. To ensure *par6^Δ226^* mutant cells were viable, we used Caspase 1 staining (Supplementary Figure 2A-B). Consistent with previous studies in the follicular epithelium (Kim et al., 2009), we found that in the absence of *par6*, aPKC and Crb were not detected at the plasma membrane of the pupal photoreceptor (Figure 2A-B). Instead of being detected at the apical membrane, Crb staining showed punctate structures in the apical region of the cytosol (Figure 2B). However, Baz and Arm domains were still present at the apical pole of the cells (Figure 2A-B) indicating that while *par6* is required for aPKC and Crb recruitment at the apical membrane, it is not required for apical-basal polarity.

**Figure 2:**
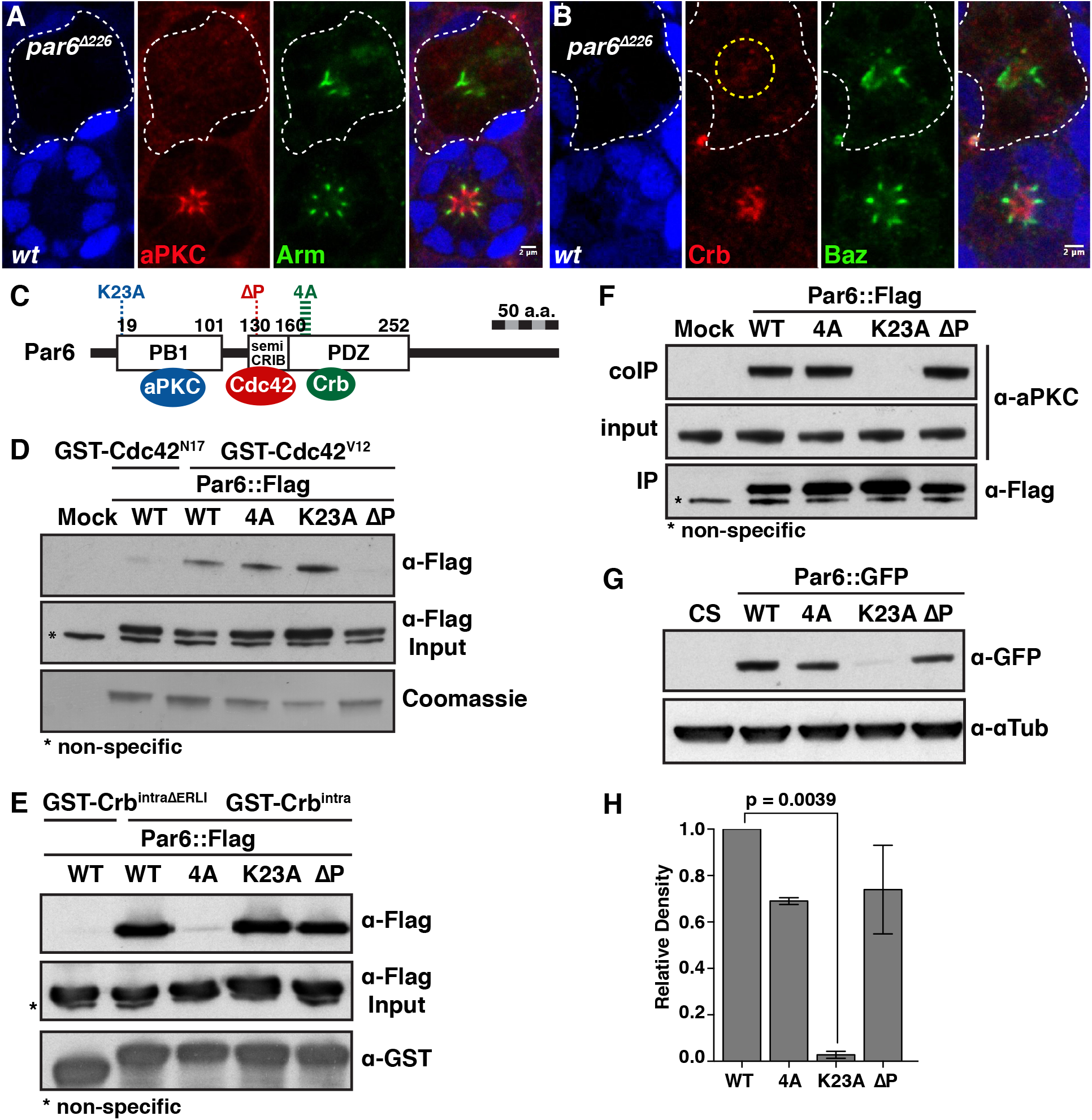
Uncoupling Par6 from Cdc42, aPKC and Crb. (**A-B**) *par6^Δ226^* mutant photoreceptors labeled by loss of GFP (blue) and stained for (**A**) aPKC (red), Arm (green) and (**B**) Crb (red), Baz (green). A dashed yellow circle indicates punctate structures, stained for Crb, in the presumptive apical region of the *par6* mutant cells. Scale bars = 2 microns. (**C**) Schematic representation of Par6, indicating the protein domains that interact with aPKC, Cdc42 and Crb, and the mutagenized sites used to uncouple these interactions. (**D**) GST-pulldown between recombinant GST::Cdc42^V12^ and SR+2 cell extracts transfected with the various *par6::Flag* transgenes. Recombinant GST::Cdc42^N17^ was used as a control. (**E**) GST-pulldown between recombinant GST::Crb^intra^ and S2R+ cell extracts transfected with the various *par6::Flag* transgenes. Recombinant GST::Crb^intraΔERLI^ (Bachmann et al., 2001) was used as a control. (**F**) Endogenous aPKC was co-immunoprecipitated from S2R+ cells transfected with the various *par6::Flag* transgenes. In (**D**) and (**F**), mock corresponds to samples transfected with empty Flag vector. (**G**) Western blot of protein extracts from adult heads of animals expressing the various *par6-Par6::GFP* transgenes, probed with anti-GFP and quantified in (**H**). Wild-type Canton S (CS) flies were used as a control. Columns represent the mean and error bars represent the SEM from 3 independent experiments.

To probe the interface between Par6, Cdc42, aPKC and Crb, we used directed mutagenesis to individually uncouple binding between Par6 and these proteins (Figure 2C). To disrupt Par6 binding to Cdc42, we used Par6^Δ139P^ (Par6^ΔP^), a mutated protein that cannot bind to Cdc42 (Hutterer et al., 2004). Interrupting Par6 binding to active Cdc42 (Figure 2D) did not interfere with the ability of this protein to bind to Crb or aPKC (Figure 2E-F). To disrupt Par6 binding to Crb, we generated Par6^KPLG170-173AAAA^(Par6^4A^) (Joberty et al., 2000; Li et al., 2010; Peterson et al., 2004; Whitney et al., 2016), a protein that retained its ability to bind to Cdc42 and aPKC, but cannot bind to Crb (Figure 2D-F). Finally, to uncouple Par6 from aPKC, we generated Par6^K23A^ (Noda et al., 2003), in which Par6 binding to aPKC was abolished without affecting its binding to Cdc42 or Crb (Figure 2D-F). All Par6 transgenes were GFP-tagged and placed under the control of a minimal *par6* promoter in order to generate transgenic animals. Western blotting from *in vivo* head extracts showed that all fusion proteins, except for Par6^K23A^, could be detected and were expressed at similar levels (Figure 2G-H). Because Par6^K23A^ could be stably expressed in S2R+ cells (Figure 2D-F), our results suggest that aPKC binding to Par6 is required to stabilize Par6 *in vivo*.

### aPKC regulates the apical localization of Par6

To test the suggestion that aPKC binding to Par6 is required to stabilize Par6 *in vivo*, we examined the expression of the *par6-Par6^K23A^::GFP* transgene in the retina. We found that Par6^K23A^::GFP failed to rescue the *par6*^Δ226^ phenotype (Supplementary Figure 2C) and was only detected at very low levels at the *ZA*, when expressed in otherwise wild-type cells (Supplementary Figure 2D-E). To complement this analysis, we made use of the *aPKC^psu69^* allele, which encodes a version of aPKC that does not bind to Par6 (Kim et al., 2009). In *aPKC^psu69^* mutant cells, only very low levels of aPKC were detected at the *ZA* associated with Arm (Supplementary Figure 2F and quantified in 2J). Similarly, Par6 did not accumulate at the apical membrane and instead was also found at low levels in the *ZA* (Supplementary Figure 2G-H and quantified in 2J’). Further, Par6 localization at the *ZA* was dependent on *baz* (Supplementary Figure 2K). In addition, Crb levels were lower than in wild-type cells (Supplementary Figure 2I and quantified in 2J’’). Altogether, these results show that Par6 binding to aPKC is required to promote the apical localization and accumulation of Par6 and aPKC, as well as that of Crb. However, it is not required for the recruitment of Par6 and aPKC at the plasma membrane. Our results also suggest that Par6 and aPKC each can be recruited to the *ZA* by Baz.

### Cdc42 regulates the apical localization of Par6

To better understand how Cdc42 binding to Par6 regulates epithelial morphogenesis, we made use of the *par6-Par6^ΔP^::GFP* transgene. Firstly, we asked whether Par6^ΔP^ could rescue the *par6* loss-of-function. Re-introducing Par6::GFP in *par6*^Δ226^ cells fully rescued the *par6* mutant phenotype (Figure 3A and Supplementary Figure 3A). However, this was not the case for Par6^ΔP^::GFP. Nevertheless, we detected this mutant protein toward the apical pole of the cell, where it co-localized with Baz and Arm (Figure 3B-C). In these domains, aPKC was barely detectable. In these cells, we also found that the recruitment of Par6^ΔP^::GFP required *baz* (Figure 3D). This finding is consistent with Par6 binding to Baz (Renschler et al., 2018), and with a requirement for Baz to load Par6-aPKC at the membrane.

**Figure 3:**
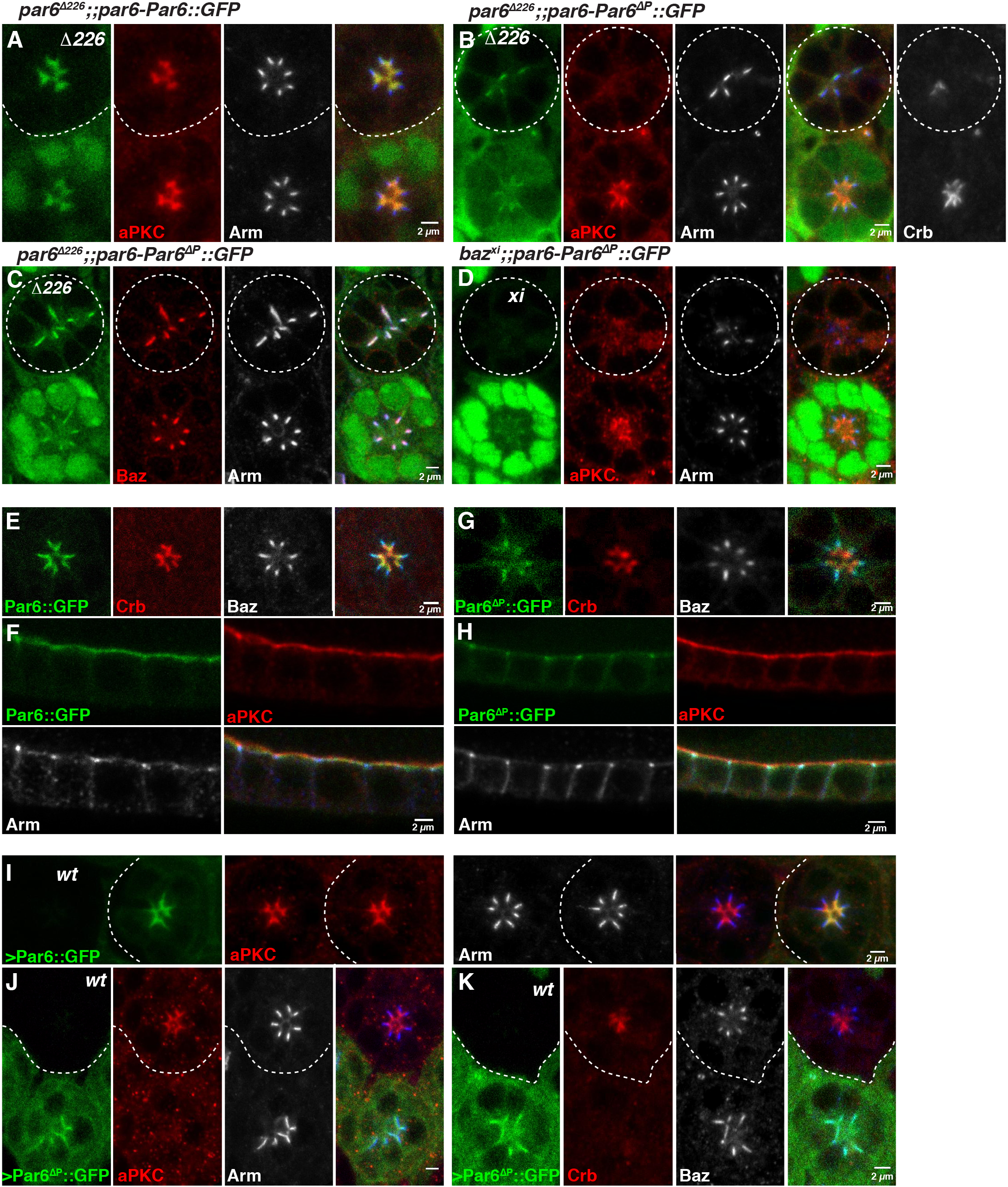
Cdc42 promotes localization of Par6 and aPKC to the apical membrane. (**A**) *par6^Δ226^* pupal retinal clones expressing *par6-Par6::GFP* (green). *par6^Δ226^* mutant cells labeled by loss of nuclear GFP signal and stained for aPKC (red) and Arm (grey). (**B-C**) *par6^Δ226^* pupal retinal clones expressing *par6-Par6^ΔP^::GFP* (green). *par6 ^δ226^* mutant cells labeled by loss of nuclear GFP signal and stained for (**B**) aPKC (red), Arm (grey); and (**C**) Baz (red), Arm (grey). In (**B**) an additional panel showing Crb staining (grey, and not part of the merged panel) is included. (**D**) *baz^xi^* clones expressing *par6-Par6^ΔP^::GFP* (green). *baz^xi^* mutant cells are labeled by loss of nuclear GFP signal and stained for aPKC (red) and Arm (grey). (**E-F**) *par6-Par6::GFP* (green) expressed in otherwise wild-type (**E**) pupal photoreceptors (Crb, red and Baz, grey) and (**F**) follicular epithelial cells (aPKC, red, and Arm, grey). (**G-H**) *par6-Par6^ΔP^::GFP* expressed in otherwise wild-type (**G**) pupal photoreceptors (Crb, red and Baz, grey) and (**H**) follicular epithelial cells (aPKC, red and Arm, grey). (**I-K**) Overexpression of wild-type Par6::GFP (**I**) and Par6^ΔP^::GFP (**J-K**) using the CoinFLP system, stained for (**I-J**) aPKC (red) and Arm (grey), (**K**) Crb (red) and Baz (grey). In all panels, the grey channel is shown in blue in the merge. Scale bars = 2 microns.

To complement these experiments, we made use of another *par6* allele, *par6^29VV^*. Using this allele we confirmed that aPKC was hardly detectable in the Baz/Par6^ΔP^::GFP domains (Supplementary Figure 3B). However, we noted instances where aPKC staining could be detected immediately apical to domains containing both Baz and Par6^ΔP^::GFP, with both *par6* alleles (Supplementary Figure 3B). Altogether, these results suggest that Cdc42 binding to Par6 regulates aPKC recruitment through Baz, and thus Par-complex assembly.

Next, to further assess the effect of uncoupling Par6 from Cdc42, we expressed the *par6-Par6^ΔP^::GFP* transgene in otherwise wild-type cells. In the photoreceptor and follicular epithelium, Par6::GFP localized as endogenous Par6 (Figure 1C, G, Figure 3E, F and Supplementary Figure 3D-E, H). In contrast, Par6^ΔP^::GFP was found at low levels at the developing *ZA* (Figure 3G, H and Supplementary Figure 3F-G, I). Furthermore, failure of Par6^ΔP^::GFP to localize at the apical membrane occurred even though Crb was present (Figure 3G and Supplementary Figure 3G and I). Quantification of the intensity levels also suggested that the interaction between Par6 and Cdc42 is required for the accumulation of Par6 at the plasma membrane (Supplementary Figure 3J).

To further probe the relationship between Par6 and aPKC, Crb, we also overexpressed Par6^ΔP^::GFP using the UASp-Par6^ΔP^::GFP transgene. While overexpression of wild type Par6::GFP had no discernable effect (Figure 3I), overexpression of Par6^ΔP^::GFP led to a strong decrease in the apical levels of aPKC and Crb when compared to the neighboring wild-type cells, and Par6^ΔP^::GFP localization was restricted to domains containing Arm and Baz, and to the cytosol (Figure 3J-K). These effects on aPKC and Crb recruitment likely reflect Par6^ΔP^::GFP outcompeting endogenous Par6 at the plasma membrane. They confirm that Cdc42 binding to Par6 regulates the apical recruitment of aPKC, and are compatible with the hypothesis that together, Cdc42, Par6 and aPKC promote the apical accumulation of Crb.

### Crb promotes the retention of Par6-aPKC at the apical membrane

To further probe the relationship between Crb and Par6, we generated *crb^11A22^* mutant cells. In these cells, the Baz and Arm domains are longer that in wild-type cells (Figure 4A-B) (Walther and Pichaud, 2010). Nevertheless, a reproducible apical fraction of Par6 and aPKC was separated from Baz and Arm. Quantification of the Par6 and aPKC signals along the apical-basal axis showed that their amount at the apical membrane was lower than in wild-type cells and these two proteins spread towards the basal pole of the cells (Figure 4C-E). Further analysis revealed that the apical membranes labeled by Par6 and aPKC were longer than those measured in wild-type and the *ZA* was wider (Figure 4E-F). Because we found Cdc42 is present at the apical membrane (Figure 1K-L,N) where Baz levels are very low (Figure 1C), we hypothesize that the apical Par6-aPKC fraction detected in *crb^11A22^* mutant cells corresponds to Cdc42-Par6-aPKC.

**Figure 4:**
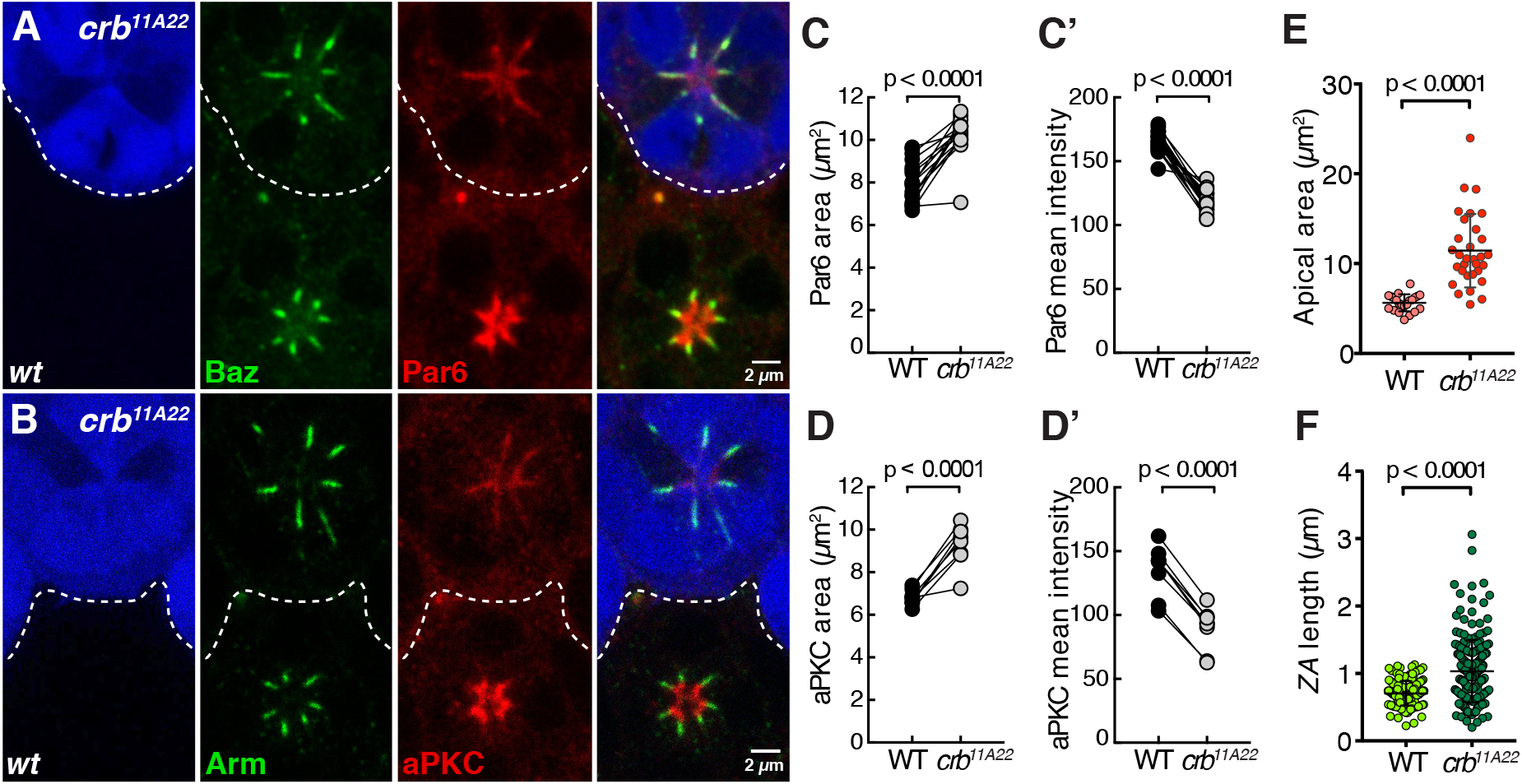
Crb is required for depleting Par6-aPKC from the *ZA*. (**A-B**) *crb^11A22^* mutant pupal photoreceptors positively labeled by GFP (blue) and stained for (**A**) Baz (green) and Par6 (red), (**B**) Arm (green) and aPKC (red). Scale bars = 2 microns. (**C-D**) Par6 and aPKC area and mean intensity quantifications in *crb^11A22^* mutant photoreceptors compared to wild-type. 20 ommatidia pairs from 7 retina and 9 ommatidia pairs from 3 retinas were measured for Par6 and aPKC, respectively. (**E**) Apical areas measured in wild-type and *crb^11A22^* mutant photoreceptors (**F**) *ZA* lengths measured in wild-type and *crb^11A22^* mutant photoreceptors.

Next, to better understand the relative contribution of Par6 binding to Crb we made use of the *par6-Par6^4A^::GFP* transgene. When expressed in otherwise wild-type cells, the apical levels of Par6^4A^::GFP were lower than that measured for Par6::GFP (Supplementary Figure 4A-C). These results support our model that Par6 binding to Crb is required for the accumulation of Par6 at the apical membrane. In addition, we observed a basal displacement of the Par6^4A^::GFP signal, as this protein was detected at the lateral membrane of the pupal photoreceptor and follicular epithelium (Figure 5A-D; quantified in 5E-F’). These results show that one contribution of Par6 binding to Crb is to promote the apical retention and accumulation of Par6 and aPKC at the developing apical membrane.

**Figure 5:**
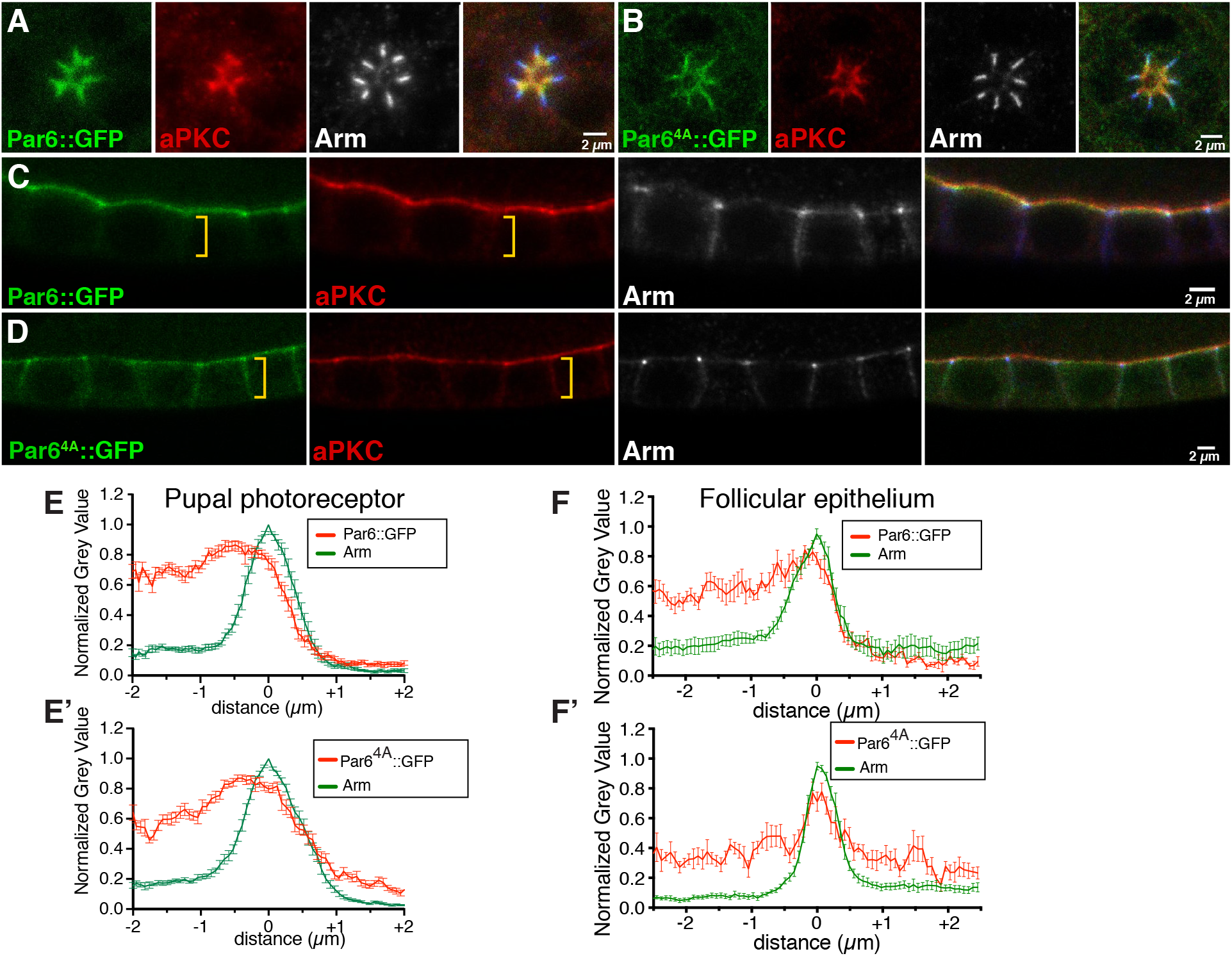
Crb promotes apical retention of Par6 and aPKC. (**A**) Expression of *par6-Par6::GFP* (green) and (**B**) *par6-Par6::GFP* (green) in otherwise wild-type photoreceptors and stained for aPKC (red) and Arm (grey). (**C**) *par6-Par6::GFP* (green) and (**D**) *par6-Par6^4A^::GFP* (green) expressed in otherwise wild-type follicular epithelial cells stained for aPKC (red) and Arm (grey). A yellow bracket indicates the lateral membrane. In all panels, the grey channel is shown in blue in the merge. Scale bars = 2 microns. (**E – E’**) Intensity profiles of Arm and Par6 at the cell cortex and plasma membrane from the apical domain towards the lateral domain of wild-type pupal photoreceptor expressing (**E**) *par6-Par6::GFP* or (E’) *par6-Par6^4A^::GFP*. Lines represent the mean value of 12 photoreceptors from 3 independent retinas, error bars show the SEM. (**F – F’**) Intensity profiles of Arm and Par6 at the cell cortex and plasma membrane from the apical domain towards the lateral domain of wild-type follicle cells expressing (**F**) *par6-Par6::GFP* or (**F’**) *par6-Parβŕ^A^::GFP*. Lines represent the mean value and error bars show the SEM (n = 6 cells from 2 individuals for *par6-Par6::GFP* and n = 9 from 3 individuals for *par6-Par6Γ^A^::GFP)*. In E-F’, the highest intensity measurement at the ZA defines the 0 value on the (x) axis.

We also assessed the ability of Par6^4A^::GFP to rescue the apical morphogenesis phenotype of *par6^Δ226^* mutant cells. These experiments showed that Par6^4A^::GFP cannot support the apical accumulation of aPKC and Crb (Figure 6A-B and quantified in 6C-C’). In these cells, Par6^4A^::GFP was detected at the apical pole of the cells, with low levels at the apical membrane and higher levels in Arm-(and PS980-Baz) labeled *AJ* domains (Figure 6B,D). Finally, when overexpressed, Par6^4A^::GFP was detected at the apical membrane and *ZA*, as well as in the cytosol (Supplementary Figure 4D). In this case, apical accumulation of aPKC was reduced (Supplementary Figure 4D-E). These results are consistent with Par6^4A^::GFP outcompeting Par6. Overall, from these experiments we conclude that Par6 binding to Crb promotes Par6 and aPKC apical retention and accumulation.

**Figure 6:**
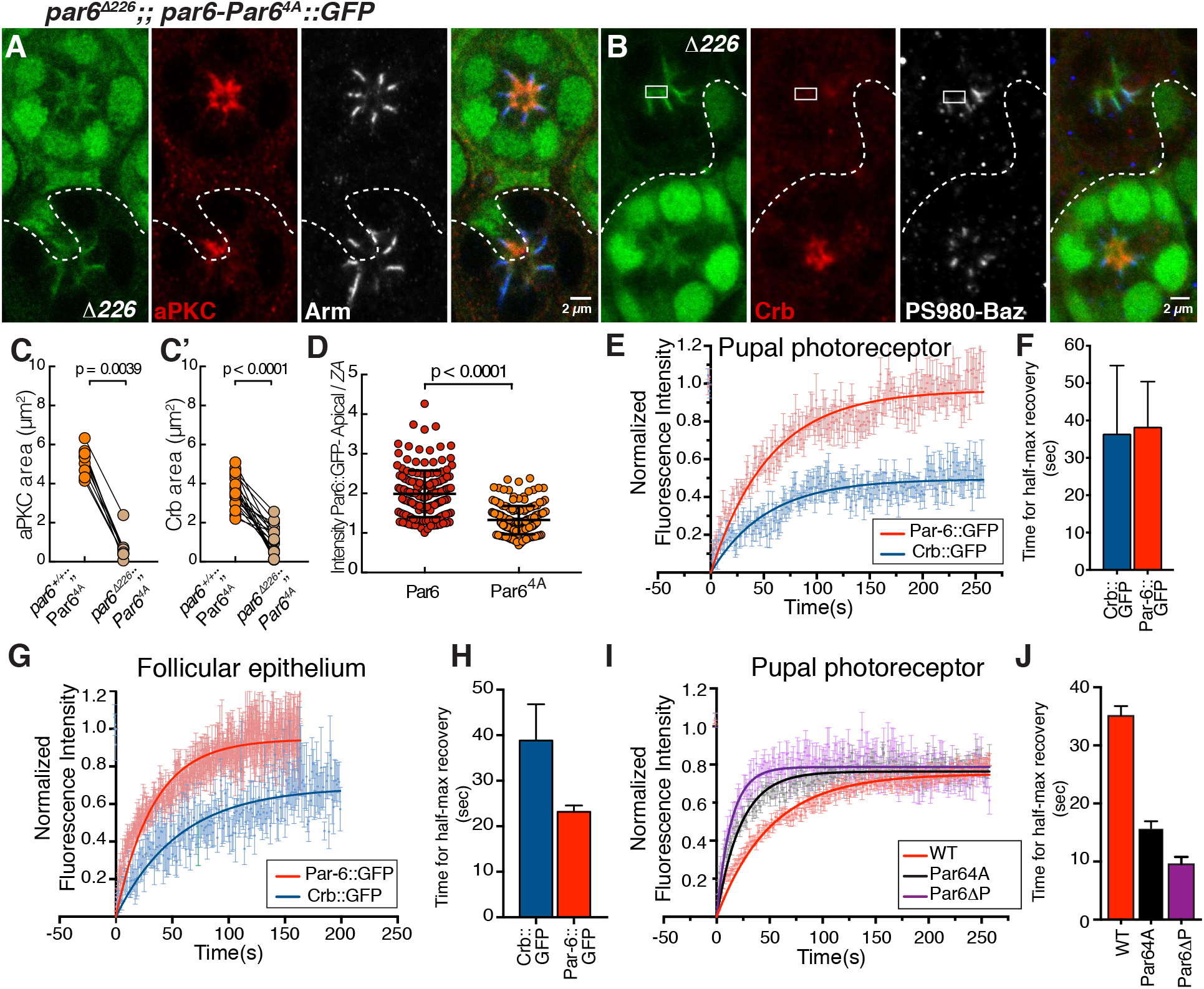
Crb stabilizes Par6 at the apical membrane. (**A – B**) *par6^Δ226^* cells expressing *par6-Par6^4A^::GFP* (green). *par6^Δ226^* mutant cells labeled by loss of nuclear GFP signal and stained for (**A**) aPKC (red), Arm (grey), (**B**) Crb (red), P-S980Baz (grey). A rectangle highlights *par6-Par6^4A^::GFP* signal detected immediately apical to P-S980Baz (C-C’) Quantification of aPKC and Crb area in the rescue experiments. 9 ommatidia pairs from 4 retinas and 19 ommatidia pairs from 6 retinas were analyzed for aPKC and Crb, respectively. (D) Quantification of the relative distribution of Par6::GFP and Par6^4A^::GFP between the apical membrane and *ZA*. At least 160 ratios were calculated from at least 3 retinas per genotype. (**EH**) FRAP of Crb::GFP (blue) and Par6::GFP (red) in the developing photoreceptor (**E**), and the follicular epithelia (**G**). Fluorescence recovery curves were calculated using a single exponential fit of the FRAP data. The half-time recoveries of Crb::GFP (blue) and Par6::GFP in the photoreceptor and follicular epithelia are shown in (**F**) and (**H**) respectively. (**I**) FRAP of Par6::GFP, Par6^4A^::GFP and Par6^ΔP^::GFP overexpressed using the Gal4-UAS system. The graph shows mean normalized fluorescence intensity for Par6::GFP (red, n = 14 from 3 individuals), Par6^4A^::GFP (grey, n = 16 from 3 individuals) and Par6^ΔP^::GFP (purple, n = 13 from 2 individuals). Error bars represent SEM. Fluorescence recovery curves were calculated using a single exponential fit. (**J**) Half-time recovery of Par6::GFP (red), Par6^4A^::GFP (black) and Par6^ΔP^::GFP (purple).

### Crb stabilizes Par6 at the apical membrane

One possibility is that Par6 binding to Crb stabilizes Par6. To test this, we used fluorescence recovery after photobleaching (FRAP) and assessed the recovery rate of Par6 and Par6^4A^::GFP in pupal photoreceptors and follicle cells. FRAP experiments of endogenously tagged Par6 over approximately 4 minutes showed that up to 95% of Par6 is mobile, with a t_1/2_ of ~40 seconds in the photoreceptor (Figure 6E-F). We found a similarly high mobile fraction for Par6::GFP in the follicular epithelium, but the estimated t_1/2_ of ~25 seconds was shorter than that measured in the retina (Figure 6G-H). These results show that overall most of the Par6 is mobile over a period of 4 min in both epithelia. The estimated t_1/2_ times also indicate that Par6 turns over more rapidly in the follicular epithelium relative to the pupal photoreceptor, suggesting that different regulations might take place at the apical membrane of these two cell types.

Expression of Par6^4A^ under the minimal *par6* promoter leads to a weak apical accumulation (Supplementary Figure 4A-C), therefore limiting our FRAP of this construct to overexpression experiments. When overexpressed in the pupal photoreceptor, Par6 and Par6^4A^ showed comparable mobile fractions (~80%). However, Par6^4A^ recovered twice as fast as Par6, with a t_1/2_ of ~ 35 seconds for UAS-Par6::GFP compared to a t_1/2_ of ~ 15 seconds for UAS-Par6^4A^::GFP (Figure 6I-J). These results suggest that Crb can either limit the lateral diffusion of Par6 or regulate its on/off rate at the apical membrane. In parallel, we estimated the mobile fraction and t_1/2_for the fraction of Par6^ΔP^::GFP associated with *AJ* material when overexpressed in the pupal photoreceptors. We found that the mobile fraction of this protein is similar to that of Par6 and Par6^4A^::GFP (Figure 6I). We estimated the t_1/2_ of UAS-Par6^ΔP^::GFP to be ~ 10 seconds (Figure 6J). This indicates that a version of Par6 that does not bind to Cdc42 (Par6^ΔP^::GFP) is more dynamic than a version that does not bind to Crb (Par6^4A^::GFP) and Par6::GFP.

To complement this analysis, we also performed Crb::GFP FRAP. In the pupal photoreceptor, we found that over a period of ~ 4 minutes, approximately 40% of Crb is mobile (Figure 6E). The mobile fraction recovers with a t_1/2_ of ~ 40 seconds, a value that is similar to that estimated for the Par6 mobile fraction over the same time span (Figure 6F). In the follicular epithelium, we found that up to 60% of Crb::GFP is mobile, with a t_1/2_ of ~ 40 seconds (Figure 6G-H). Therefore while the rate of recovery for Crb is similar in both epithelia, a marginally larger fraction is mobile at the apical membrane of the follicular epithelial cells when compared to the pupal photoreceptor. From these results, we conclude that different regulations might take place in the follicular epithelium and pupal photoreceptor when considering the dynamics of Par6 and Crb at the apical membrane. Overall, a greater fraction of Crb is stable at the apical membrane of these two epithelial cell types when compared to Par6, pointing to a dynamic relationship between these two proteins.

## DISCUSSION

How Cdc42, the Par-complex (Baz/Par3 and Par6-aPKC) and Crb/CRB3 come together to regulate epithelial morphogenesis is not well understood. With this study, we present evidence that Cdc42 is active at the apical membrane and *ZA* of the pupal photoreceptor, and show that Cdc42 binding to Par6 promotes apical localization of Par6-aPKC. In addition, we find that Cdc42 promotes the apical retention of Par6-aPKC through Crb, and this retention mechanism depends on Par6 binding to Crb. Our work in the developing follicular epithelium indicates that similar regulations take place in these cells.

### Cdc42 and aPKC regulate Par6 apical localization

We find that in both the pupal photoreceptor and follicular epithelium, Par6 binding to Cdc42 is required for Par6 to be localized at the apical membrane. In rescue experiments in the pupal photoreceptor, we find that the corresponding Par6ΔP protein colocalizes with *AJ* material. Further, its expression in otherwise wild-type cells leads to its localization at the *ZA*, and this localization depends on *baz*. These findings differ from the situation reported in the embryonic ectoderm, where a similar (untagged) version of Par6 failed to be recruited at the membrane altogether (Hutterer et al., 2004). They also raise the possibility that during epithelial morphogenesis, Par6-aPKC are recruited to the *ZA* by Baz, prior to being captured by Crb. Furthermore, our finding that Par6 and aPKC both localize at the *ZA* when binding between these two proteins has been interrupted genetically using the *aPKC^psu69^* allele, raises the possibility that Baz can recruit these two proteins independently. This conclusion is consistent with the observation that in the cellularizing embryo, Baz supports aPKC localization independently of Par6 (Harris and Peifer, 2005).

### Cdc42 promotes Par6-aPKC retention through Crb

Our recent analysis of *baz* mutant pupal photoreceptors suggests that Baz is required to load Par6-aPKC at the apical membrane, however low levels of these proteins can still be detected in a fraction of *baz* mutant cells (Walther et al., 2016). These low levels of Par6-aPKC indicate that they can be recruited independently of Baz, presumably through binding to Cdc42 or Crb. Here, we show that Par6-aPKC can be detected at the apical membrane of *crb* mutant cells where we do not detect Baz. In addition, our rescue experiments expressing the version of Par6 that cannot bind to Crb show that Par6-aPKC can localize apical to *AJ* domains. Altogether, we hypothesize that this apical fraction of Par6-aPKC is bound to Cdc42 and therefore that both Cdc42 and Baz are able to support apical recruitment of Par6 and aPKC during epithelial morphogenesis. The lack of a requirement for Baz in the follicular epithelium (Shahab et al., 2015) could reflect a more preeminent role for Cdc42 in loading Par6-aPKC at the membrane of these cells when compared to other epithelia.

Common to the pupal photoreceptor and follicular epithelium, we find that while Crb is not required for localizing Par6-aPKC at the apical membrane, it is required to promote their apical accumulation through retention. This conclusion is well supported by our FRAP experiments showing that the t_1/2_ recovery of the Par6^4A^ mobile fraction is half of that of wild type Par6. Further, our FRAP experiments indicate that over 4 minutes, up to 95% of Par6 is mobile at the apical membrane. This is in contrast to Crb, as we find that in the pupal photoreceptor, approximately 40% is mobile over the same time span. This difference is likely to reflect that the Crb-Par6 interaction is dynamic and also that Crb turnover depends at least in part on endocytosis and recycling, which might operate at longer time scales (Lin et al., 2015; Perez-Mockus et al., 2017). We also find that when compared to the pupal photoreceptor, in the follicular epithelium a greater proportion of Crb is mobile (60%), indicating that differences exist between these two tissues when considering the relative stability of Crb. Nevertheless, our results indicate that in both cell types Cdc42 promotes Par6 binding to Crb to stabilize and thus accumulate Par6-aPKC at the apical membrane. Reciprocally, Par6-aPKC promotes the accumulation of Crb. Therefore our work indicates that during epithelial morphogenesis, Cdc42 promotes apical identity by coordinating the apical recruitment of Par6-aPKC and Crb.

## Acknowledgements

We thank A. Wodarz, J. Knoblich, F. Wirtz-Peitz, D. St Johnston, E. Knust, B. Thompson, B.Baum, N. Tapon, and M. Meitzstein for providing reagents. We thank the Pichaud lab members for discussion and support, and in particular Katherine Massey her contribution in supervising M.P. The N2 A71 anti-Armadillo, DCAD2 anti-Ecad and Cq4 anti-Crb antibodies, developed by E. Weischaus, T. Uemura and E. Knust respectively, and were obtained from the Developmental Studies Hybridoma Bank, created by the NICHD of the NIH and maintained at The University of Iowa, Department of Biology, Iowa City, IA 52242. Stocks obtained from the Bloomington Drosophila Stock Center (NIH P40OD018537) and the Vienna Drosophila Resource Center were used in this study. This work was funded by an MRC grant to FP (MC_UU_12018/3).

## Author contribution

F.P. and R.F.W. supervised the work and conceived the project together with F.N.A. R.F.W performed the mutagenesis and cloned all *par6* transgenes; conducted and analyzed all Par6 related experiments in the pupal photoreceptor; performed and analyzed the FRAP experiments in the pupal photoreceptor. F.N.A. performed and analyzed all biochemical experiments; performed and analyzed all the experiments in the follicular epithelium, including FRAP of Par6 and Crb. E.V. and M.P. contributed to the cloning. F.P. wrote the manuscript together with F.N.A. and R.F.W.

## MATERIAL AND METHODS

### Fly strains

The following fly strains were used:

*par6^Δ226^, FRT9.2* (Petronczki and Knoblich, 2001)
*par6^29VV^, FRT19A* (Jones and Metzstein, 2011)
*w, baz^xi106^, FRT9.2* (Nusslein-Volhard et al., 1987)
*w; FRTG13, aPKC^psu69^* (Kim et al., 2009)
*w;; FRT82B, crb^11A22^* (Tepass et al., 1990)
*w; (sqh-ChFP-Cdc42)^23^* and *w;; (sqh-ChFP-Cdc42)^33^* (Abreu-Blanco et al., 2014)
*w, cdc42^3^, FRT9.2* and *w, cdc42^4^, FRT19A* (Fehon et al., 1997)
*w; cdc42^IR^* [Vienna *Drosophila* Resource Center (VDRC) 100794]
*w;; baz^IR^* [Bloomington *Drosophila* Stock Center (BDSC) 35002]
*w* ; *GMR-Gal4* (Freeman, 1996).
*w; GR1-Gal4* (Goentoro et al., 2006)

The following fly strains were generated in this study: *w* ;; *par6-Par6::GFP, w ;; par6-Par6^K23A^::GFP, w* ;; *par6-Par6^ΔP^::GFP, w* ;; *par6-Par6 ^4A^::GFP, w;; UASp-Par6::GFP, w;; UASp-Par6^4A^::GFP, w;; UASp-Par6^ΔP^::GFP. UASp-WASp-CRIB::GFP and UASp-WASp-CRIB::GFP^MUT^*. To generate *par6-Par6::GFP* rescue strains, the appropriate DNA constructs were injected into the parent strain *y1, w67c23;; P{Cary} attP2*, [Bloomington *Drosophila* Stock Centre (BDSC) 8622] for PhiC31 mediated recombination (Groth et al., 2004) by BestGene Inc. *UASp-Par6::GFP (wild type, ΔP and 4A), UASp-WASp-CRIB::GFP and UASp-WASp-CRIB::GFP^MUT^* strains were generated by injecting the appropriate DNA constructs for standard P-element transformation (BestGene Inc.) (Rubin and Spradling, 1982).

### Molecular Biology

*par6* cDNA (Clone LD29223) was obtained from the *Drosophila* Genomics Resource Centre, Indiana and cloned into the *pENTR™/D-TOPO®* vector (Invitrogen) to generate *pENTR-Par6* cDNA. The *par6* genomic construct, *pENTR-H427* (unpublished), was a kind gift of Frederick Wirtz-Peitz (Knoblich lab). *pENTR-H427* contains a 1kb minimal *par6* promoter (~1kb region upstream of the ATG start codon), the *par6* coding region with a C-terminal GFP fusion and a minimal *par6* 3’UTR. Following sequence verification of both *par6* pENTR starting vectors, the QuikChange Mutagenesis System (Agilent) was used to generate the Par6^K23A^, Par6^ΔP^ and Par6^4A^ substitutions. Par6^ΔP^ is a deletion of P139 while the Par6^4A^ is the substitution of residues KPLG170-173 for Alanines. The mutated *par6* pENTR vectors were sequence verified and used for cloning with the Gateway™ Cloning system (Invitrogen). The destination vector *pBID-G* (Addgene 35195) was used in combination with *pENTR-H427* derived mutants to generate all *par6*-Par6::GFP rescue constructs. All other *par6* DNA constructs used in this study are derivatives of *pENTR-Par6 cDNA*. The destination vector *pPWG* from the *Drosophila* Gateway™ Vector collection was used to generate *UASp-Par6::GFP, UASp-Par6^δp^::GFP* and *UASp-Par6^4A^::GFP* DNA constructs. For the *UASp-WASp-CRIB::GFP* and *UASp-WASp-CRIB::GFP^MUT^*, the CRIB domain of *Drosophila* WASp spanning residues K226 to A313 was PCR amplified from *UASp-WASp::GFP* (a gift from Buzz Baum) and cloned into the *pENTR™/D-TOPO®* vector (Invitrogen). To disrupt binding of the WASp CRIB domain to Cdc42-GTP, substitutions F240D H242D and H245D were introduced (after (Tskvitaria-Fuller et al., 2006)) using the QuikChange Mutagenesis System (Agilent). Following sequence verification of both constructs, the destination vector *pPWG* from the *Drosophila* Gateway™ Vector collection was used to generate *UASp-WASp-CRIB::GFP* and *UAS-WASp-CRIB::GFP^MUT^*.

To generate *pActin-Par6::FUAG* constructs for expression in S2R+ cells, the *pAWF* destination vector was used. *pAWF* was derived from the *Drosophila* Gateway™ Vector collection by Yanxiang Zhou (Tapon lab). cDNA constructs encoding *Cdc42^N17^* and *Cdc42^V12^* were generated and cloned into *pENTR™/D-TOPO®* vector (Invitrogen). The *pDEST15* vector, containing an N-terminal GST tag, was used to generate plasmids *GST::Cdc42^N17^* and *GST::Cdc42^N17^*using the Gateway™ Cloning System (Invitrogen). The *GST::Crb^ıntra^* and *GST::Crb^ıntraΔERU^*constructs were provided by E. Knust (Kempkens et al., 2006).

### Genetics

Clonal analysis of mutant alleles in the retina was performed using either the standard FLP-FRT technique (Xu and Rubin, 1993) with appropriate *FRT, ubi-GFP* chromosomes used to generate negatively marked mutant tissue, or using MARCM (Lee and Luo, 2001) to generate positively marked mutant tissue. In both cases, eyFLP (Newsome et al., 2000) was used. MARCM was used to generate retinal tissue expressing *baz^IR^* in the *aPKC^psu69^* mutant background. Rescue experiments with *par6-Par6::GFP* transgenes were performed by building fly strains carrying a specified mutant chromosome and the *par6-Par6::GFP* transgene of interest, followed by FLP-FRT induction of clones. Retinal clones overexpressing UAS-Par6::GFP transgenes, UAS-WASp-CRIB::GFP or UAS-WASp-CRIB::GFP^MUT^ were generated with the coinFLP system (Bosch et al., 2015) using BDSC stock 58750. UASp-Par6 transgenes were expressed in the follicular epithelium using the GR1-Gal4 driver.

### Immunofluorescence

Whole mount retinas at 40 % after puparium formation (APF) were prepared as previously described (Walther and Pichaud, 2006). Samples were fixed in 4 % Formaldehyde in PBS for 20 min, blocked in 5 % goat serum in PBS with 0.3 % Triton (PBST) for 20 min. All subsequent incubations were performed in PBST. Samples were incubated overnight at 4°C in primary antibodies, washed 3×5 minutes, incubated in secondary antibodies for 4-6 hours and washed overnight at 4 °C. Ovaries were dissected in PBS, fixed in 4 % Formaldehyde in PBS for 20 min and blocked in 5 % goat serum in PBS with 0.1 % Tween (PBT) for 20 min. For Crb stainings, ovaries were fixed in 4 % Formaldehyde in PBS for 20 min, incubated 2 min in 50 % methanol in PBT, 2 min in 100 % methanol, 2 min in 50 % methanol in PBT, washed 3×10 min in PBT and blocked 30 min in 10 % BSA in PBT. Ovaries were incubated with primary antibodies diluted in PBT overnight at 4 °C, washed 4×5 min in PBT, incubated with secondary antibodies diluted in PBT for 3h and washed 3×10 min in PBT. All incubations were at room temperature unless otherwise stated. The following antibodies were used for indirect immunofluorescence: rabbit anti-PKCζ 1/600 (SAB4502380, Sigma), mouse anti-Arm 1/200 (N27-A1, Developmental Studies Hybridoma Bank), rabbit anti-Baz 1/2000 (gift from Andreas Wodarz), rat anti-Crb 1/200 (Walther et al., 2016), mouse anti-Crb 1/50 (Cq4, Developmental Studies Hybridoma Bank), guinea pig anti-Par6 1/400 (Walther et al., 2016), rat Ecad 1/20 (DCAD2, Developmental Studies Hybridoma Bank), rabbit anti-Sdt 1/400 (gift from E. Knust), with the appropriate combination of mouse, guinea pig, rabbit and rat secondary antibodies conjugated to Dy405, Alexa488, Cy3 or Cy5 as appropriate at 1/200 each (Jackson ImmunoResearch). Samples were mounted in VectaShield™ with or without DAPI as appropriate and imaging was performed using a Leica SP5 or SP8 confocal microscope. Images were edited using Fiji and Adobe Photoshop 7.0.

### Quantifications

To measure pixel intensity and area of epitope staining a threshold was applied to define the domain(s) of interest and then quantified using the wand (tracing) tool in Fiji. A minimum of 12 data points were obtained for each condition, from at least 3 independent retinas. To estimate relative protein distribution between the apical membrane and *ZA* at least 60 ratios were measured from at 3 independent retinas. GraphPad Prism version 7.0 for Mac was used for the statistical analyses. Data sets were tested for normality using the D’Agostino and Pearson normality test. p-values were calculated using either the student’s t-test or the Mann-Whitney test in cases where the data were unpaired, or either the paired t-test or the Wilcoxon test in cases where the data were paired.

### FRAP

FRAP analyses in the fly retinas was performed at 40% APF as previously described (Walther et al., 2016). Live imaging was performed on a Leica SP5 confocal using a 63x 1.4 NA oil immersion objective at the following settings: pixel resolution 512 x 512, speed 400 Hz, 15 % 488 nm laser power at 20 % argon laser intensity and 5x zoom. FRAP analysis of Crb::GFP (Huang et al., 2009), par6-Par6::GFP and GMR-Gal4 ; UAS-Par6 wildtype, 4A and ΔP transgenes were performed through a 5 pixel-diameter circle ROI followed by photo-bleaching with 2 pulses using 90 % 488 nm laser power at 20 % argon laser intensity. GFP recovery was recorded every 1.293 sec with the previously mentioned settings for approximately 300 sec.

FRAP analysis in the follicular epithelium was performed as previously described in (Prasad et al., 2007). Live imaging was performed on a Leica SP8 upright confocal using a 63x 1.4 NA oil immersion objective at the following settings: pixel resolution 512 x 256, speed 400 Hz, 20 % 488 nm laser power at 40 % argon laser intensity and 4x zoom. FRAP analysis of Par6::GFP and Crb::GFP were performed through a 1 μm x 1 μm square ROI followed by photo-bleaching with 2 pulses using 90 % 488 nm laser power at 40 % argon laser intensity. GFP recovery was recorded every 0.328 seconds for Par6::GFP or every 1 second for Crb::GFP with the previously mentioned settings for approximately 160-200 seconds.

For each experiment three different z-axis profiles were plotted: (1) from the photobleached area, (2) from an equivalent area of a neighboring non photo-bleached photoreceptor, and (3) from an equivalent area of background. The obtained data was normalized using easyFRAP (Rapsomaniki et al., 2012) and fitted to a two-phase association curve in GraphPad Prism version 7.0 for Mac (GraphPad Software, San Diego California USA, www.graphpad.com). Each data point represents the mean and error bars the SEM. Half-time values were determined with Prism based on the fitting curves obtained and columns represent the mean and error bars the 95 % CI of each data set. The p values were calculated with a two-way ANOVA test with Bonferoni’s correction.

### Biochemistry

To express GST-fusion proteins, *E. coli* BL21 was transformed with the appropriate plasmids and induced with 0.2 % L-arabinose or 1 mM IPTG as appropriate for 4h at 30 °C. Bacteria were lysed by sonication in 50 mM Tris HCl pH7.6, 50 mM NaCl, 5 mM MgCl_2_, 0.5 % Triton X-100, 10 mM DTT in the presence of protease inhibitor (EDTA-free Complete Protease Inhibitor [Roche]). GST-fusion proteins were purified using Glutathione Sepharose 4 Fast Flow beads (GE Healthcare), washed in lysis buffer and kept on beads in lysis buffer with 1 mM DTT at 4 °C. To express MBP-fusion proteins *E. coli* BL21 was transformed with the appropriate plasmids and induced with 0.3 mM IPTG for 2h at 37 °C. Bacteria were lysed in 20 mM Tris HCl pH 7.4, 200 mM NaCl, 1 mM EDTA, 1 mM DTT. MBP-fusion proteins were purified using Amylose resin (New England Biolabs), washed in lysis buffer, eluted in lysis buffer containing 10 mM Maltose, dialysed to 50 mM Tris HCl pH 7.5, 150 mM NaCl, 5 mM MgCl2, 40 % Glycerol and stored at −80 °C for further experiments.

*Drosophila* Schneider S2R+ cells (DGRC) were transiently transfected using Effectene Transfection Reagent (Qiagen) with empty vector *pActin-Flag* (Mock) or *pActin-par6-Flag* (WT, K23A, 4A or ΔP) and lysed in 50 mM Tris HCl pH 7.5, 150 mM NaCl, 0.5 % Triton X-100, 1 mM EDTA, protease inhibitor (EDTA-free Complete Protease Inhibitor [Roche]) and phosphatase inhibitor cocktail (Sigma). For co-immunoprecipitation experiments, Par6::Flag was immunoprecipitated with antiFLAG M2 magnetic beads (Sigma) for 1h at 4 °C, washed three times in lysis buffer and analyzed by Western Blot. For GST pulldown experiments, S2R+ cell lysates were added to purified GST-fusion proteins for 1h at 4 °C, washed three times in lysis buffer and analyzed by Western Blot and Coomassie staining. For *in vitro* binding assays, recombinant MBP and GST proteins were incubated for 1h 4°C in binding buffer (50 mM Tris HCl pH 7.5, 150 mM NaCl, 5 mM MgCl2, 0.5 % Triton X-100) and washed three times in binding buffer. From the same sample, 20 μL was loaded onto a gel for Coomassie staining and 5 μL was loaded onto a separate gel for Western Blot analysis.

Protein extraction was performed from eight fly heads homogenized in 30 μL of extraction buffer (125 mM NaCl, 50 mM Tris HCl pH 7.5, 5 % glycerol, 1 mM MgCl_2_, 1 mM EDTA, 0.2 % NP-40, 0.5 mM DTT, protease inhibitor (EDTA-free Complete Protease Inhibitor [Roche]) and phosphatase inhibitor cocktail [Sigma]). Samples were analyzed by Western blotting. The following antibodies were used for protein detection: anti-Flag M2 mouse 1/1,000 (F3165, Sigma), anti-myc 9E10 mouse 1/1,000 (sc-40, Santa-Cruz), anti-GFP (D5.1) XP rabbit 1/1,000 (2956S, Cell Signaling), anti-αTubulin mouse 1/100 (AA4.3, DSHB), anti-GST rabbit 1/100,000 (G7781, Sigma), anti-MBP mouse 1/80,000 (E8032S, New England Biolabs). Western Blots were quantified using Fiji, while graphical representation and statistical analysis were performed in GraphPad Prism version 7.0 for Mac. Columns represent mean, and error bars are the SEM of each dataset. p values were calculated with a Kruskal Wallis test and corrected using Dunn’s multiple comparison test.

